# Structural modelling and biophysical analyses reveal a dimeric coiled-coil architecture in the FAZ10 central region of *Trypanosoma brucei*

**DOI:** 10.64898/2026.05.05.722045

**Authors:** Cleidy Mirela Osorio-Mogollon, Diego Antonio Leonardo, Clarice Izumi, Leticia Cioca Alves, Gustavo E. Olivos-Ramirez, Adolfo B. Poma, Munira Muhammad Abdel Baqui

## Abstract

*Trypanosoma brucei* relies on the flagellum attachment zone (FAZ) to coordinate flagellum positioning, cell morphology, and cytokinesis. The giant FAZ10 protein, which contains both repetitive and structured regions, is essential for correct cleavage-furrow placement, yet its molecular organization remains unresolved due to its exceptional size. In this study, we define the architecture of the FAZ10 central region through structural modelling and biophysical validation. AlphaFold2 and molecular dynamics simulations defined a parallel coiled-coil dimer flanked by symmetric globular domains, with canonical hydrophobic core packing, complementary interhelical contacts, and local heptad discontinuities, including a stammer–stutter pair that modulates the superhelical geometry. Biophysical analyses show that this region forms a stable dimer in solution, mediated by the coiled-coil domain and consistent with a predominantly ***α***-helical structure. Together, these findings identify the FAZ10 central region as a semi-flexible dimeric scaffold that provides a structural framework for understanding FAZ supramolecular organization and the integration of large cytoskeletal assemblies in trypanosomatids.

## Introduction

Neglected tropical diseases remain a major global health concern, even as interventions declined from 2.2 billion in 2010 to 1.5 billion in 2023 [1]. As a result, the objectives of the WHO 2030 roadmap remain challenging to achieve, partly due to persistent gaps in our understanding of the molecular and structural principles underlying parasite biology and pathogenicity [2]. Human African Trypanosomiasis (HAT) or sleeping sickness exemplifies this challenge, as it continues to impose a substantial public health burden in sub-Saharan Africa [3]. HAT is caused by the unicellular flagellated parasite *Trypanosoma brucei*, which is transmitted to humans through the bite of *Glossina* spp. (tsetse flies) [4].

Many essential aspects of *T. brucei* biology are controlled by the cytoskeleton, a highly organized cellular architecture that governs cell morphology, polarity, motility, and division [5–7]. Central to this organization is the Flagellum Attachment Zone (FAZ), a multiprotein complex that anchors the flagellum along the cell body, coordinates morphogenesis and cytokinesis, and acts as a structural scaffold [8–10]. The FAZ exhibits a compartmentalized architecture comprising a flagellar domain, an intra-cellular domain, the FAZ filament, the microtubule quartet, and associated linker structures, assembled through an extensive network of protein–protein interactions [6, 11–13]. However, the molecular and structural organization of the FAZ filament scaffold remains poorly defined.

FAZ10 is a giant FAZ protein of 4,334 amino acids that migrates at approximately 2 MDa on SDS–PAGE, consistent with an extended and highly repetitive architecture [14]. It localizes to the FAZ intracellular domain, where it contributes to the staple-like structures that anchor the flagellum to the cell body [14]. Depletion of FAZ10 disrupts cleavage furrow positioning during cytokinesis, leading to premature or asymmetrical cell division and severe morphogenetic defects [14]. FAZ10 is organized into five regions, including two extensive repeat domains composed of 52 nearly identical 35-residue repeats each, flanked by N- and C-terminal segments and a central region. Despite its clear functional importance, structural information for FAZ10 is lacking. Its exceptional size and repetitive organization likely hinder conventional structure determination, motivating a reductionist focus on smaller tractable subregions. Defining the architecture of these regions is therefore essential to understand how FAZ10 contributes to the mechanical framework of the FAZ.

Here, we combine structural modelling and biophysical experiments to define the molecular arrangement of the FAZ10 central region from *T. brucei*. AlphaFold2-based predictions and coarse-grained molecular dynamics simulations using the Martini 3 framework were used to generate a structural model of the assembly and probe its conformational behaviour. In parallel, size-exclusion chromatography (SEC), SEC coupled to multi-angle light scattering (SEC–MALS), and circular dichroism (CD) spectroscopy were used to determine its oligomeric state and structural properties. Together, these analyses show that the FAZ10 central region adopts an ordered, predominantly *α*-helical dimeric assembly mediated by a long coiled-coil domain. Rather than behaving as a rigid rod, this elongated structure accommodates localized bending and limited axial flexibility, consistent with a semi-flexible filamentous scaffold. Our findings provide the first structural characterization of FAZ10 and suggest that its dimeric organization supports the mechanical architecture of the full-length protein within the FAZ cytoskeleton.

## Results

### Sequence-based predictions reveal the central organization of FAZ10

IUPred2A analysis of the full-length FAZ10 sequence reveals extensive intrinsically disordered regions (IDRs), primarily concentrated within repetitive regions, indicating a non-uniform distribution along the sequence (Fig. 1A). Thus, FAZ10 was subdivided into five regions: the N-terminal, central, and C-terminal domains, as well as two repetitive regions. Among these, the central region (D2070–P2576) exhibited markedly lower predicted disorder (26.04% on average) compared to the N-terminal (41.16%), C-terminal (28.65%), and the two repetitive regions (96.72% and 86.95%, respectively). A multiple sequence alignment of the central region across different *T. brucei* strains showed that this region is conserved in length and displays only limited sequence variation (Fig. S1), supporting its preservation both in size and sequence across strains. Based on these observations, the central region was selected for detailed structural and biophysical characterization. Secondary structure prediction using PSIPRED [15] indicated a predominance of *α*-helical content (Fig. 1B, S2). Coiled-coil probability was then evaluated using the MARCOIL algorithm [16]. These data reveal a canonical heptad repeat pattern (abcdefg) with probabilities *>*90% along the sequence (Fig. 1C). MARCOIL further predicted a consistent coiled-coil dimer organization within the central region throughout the helical wheel representation (Fig. 1D). Here, hydrophobic residues at ‘a’ and ‘d’ positions of the heptad repeat align across the interface. They are abundant and form the hydrophobic core of the coiled-coil assembly.

**Fig. 1:**
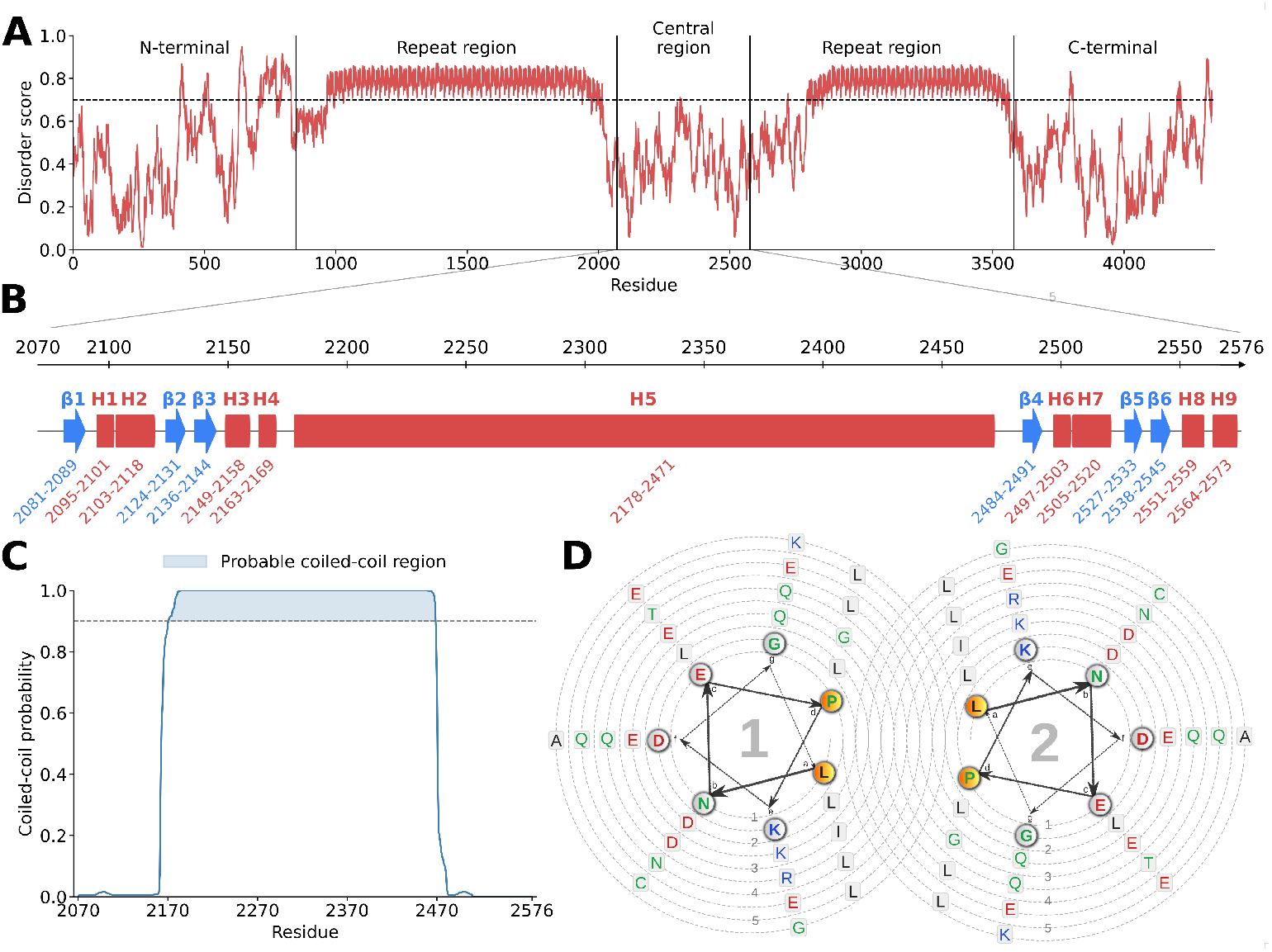
Predicted structural features of FAZ10. (A) Intrinsic disorder profile of the full-length protein 4,334–amino acid, by IUPred2A analysis showing five regions: N-terminal, central, and C-terminal domains, and repetitive regions (threshold = 0.70). (B) Secondary-structure annotation of the central region determined by PSIPRED (*β*: beta sheet, H: helix). (C) Central region was analysed by MARCOIL, confirming the probability of coiled-coil structure (threshold = 0.90). (D) Helical wheel projections of the predicted coiled-coil segment showing residue arrangement according to the canonical heptad repeat and a parallel dimeric conformation. Numbers 1–5 denote the successive radial windings of the wheel from the center outward, and arrows indicate the direction of sequence progression along the wheel spiral.

### Structural prediction of the FAZ10 central region

The central region of FAZ10 was modeled as a homodimer using AlphaFold2 [17] (Fig. S3-A). Confidence metrics (pLDDT, PAE, and sequence coverage) were high across most of the structure, indicating high confidence in the predicted model (Figs. S3-B-D). The initial model adopts an elongated architecture composed of two parallel coiled *α*-helices approximately 50.8 nm in length, flanked at both ends by globular domains. These globular domains share a similar fold, comprising a small *β*-sheet formed by three antiparallel *β*-sheets and four *α*-helices (Fig. S3-E). In general, the model forms a symmetric dimer with a twofold rotational symmetry axis parallel to the coiled-coil. Consistent with this arrangement, classical knobs-into-holes packing is observed at the sequence level (Fig. S4-A) and in the structural model, where side chains from one helix insert into complementary pockets of the opposing helix (Figs. S4-B, C).

Although the AlphaFold2 model exhibited overall high confidence metrics, the predicted structure is highly homogeneous and geometrically idealized, with a perfectly aligned and fully ordered coiled-coil arrangement (Fig. S3). Such structural regularity is unlikely to fully represent the conformational heterogeneity expected under physiological conditions. To account for this limitation, we subjected the dimeric model to all-atom molecular dynamics (AA-MD) simulations for 100 ns to assess its dynamic stability, probe interhelical flexibility, and evaluate whether the predicted coiled-coil interface remains structurally stable under physiological simulation conditions.

### High-resolution molecular dynamics simulation

Further analysis of the AA-MD simulations supported the overall stability of the elongated architecture (Figs. S5-A, B). The coiled-coil domain spans approximately 41.5 nm after 100 ns (Fig. 2A), preserving the characteristic intertwining of the two *α*-helices while remaining slightly shorter than the predicted AlphaFold2 model. Despite this modest reduction in length, the assembly maintains a continuous and well-packed coiled-coil organization. A 90° rotated view reveals a subtle bend near the central portion of the filament (Fig. 2A), indicating localized flexibility along the helical axis. The terminal globular domains remain well resolved and retain their secondary structure elements throughout the simulation (Fig. 2B).

**Fig. 2:**
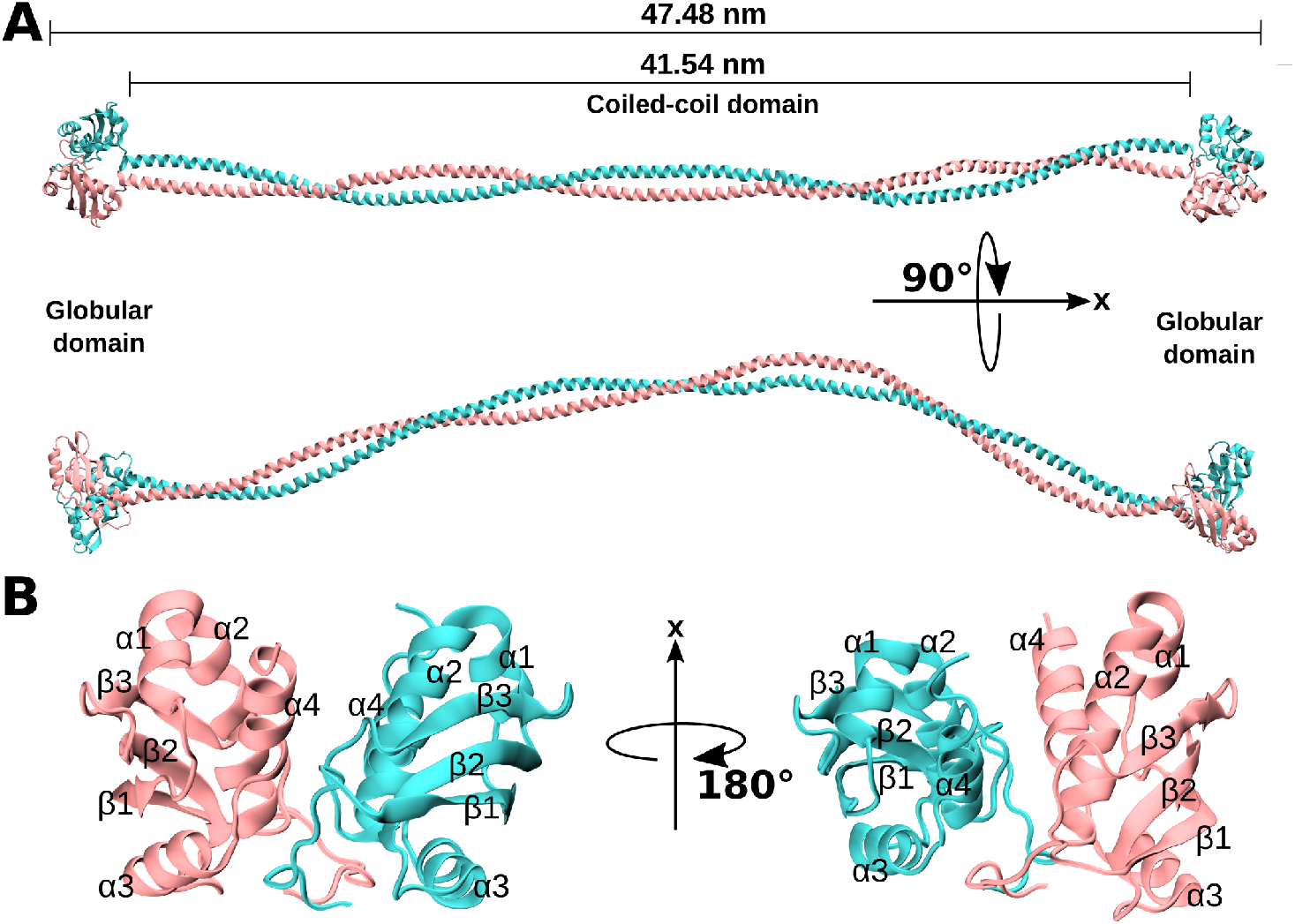
FAZ10 central region remains an elongated dimer with localized bending after atomistic molecular dynamics simulation. (A) Relaxed structure after 100 ns of all-atom MD simulation. Chains A and B are shown in pink and cyan, respectively. A 90° rotation reveals a subtle curvature along the central coiled-coil. (B) Close-up views of the globular domains at both ends of the central region.

To investigate the dynamic stability, we determined the molecular interactions that stabilize the dimer formation by analyzing the last frame of the AA-MD simulation. The coiled-coil interface is reinforced by a network of hydrophobic contacts (*e*.*g*., L2178 and L2181), salt bridges (*e*.*g*., E2247 and K2252), and hydrogen bonds (*e*.*g*., K2223 and T2224)(Figs. 3A–F), which together contribute to a tightly packed and electrostatically balanced assembly. Moreover, we observed that a salt bridge and H-bonds stabilize solvent-exposed regions of the parallel helices, further supporting structural integrity of the dimer. Furthermore, it was revealed a symmetric arrangement of residues at the ‘e’ and ‘g’ positions (Fig. 1D and 3B), with K2182 from one chain oriented toward Q2184 of the opposing chain and *vice versa* (Fig. 3C). These residues flank the hydrophobic core and form a polar belt along the dimer interface (Fig. 3C).

**Fig. 3:**
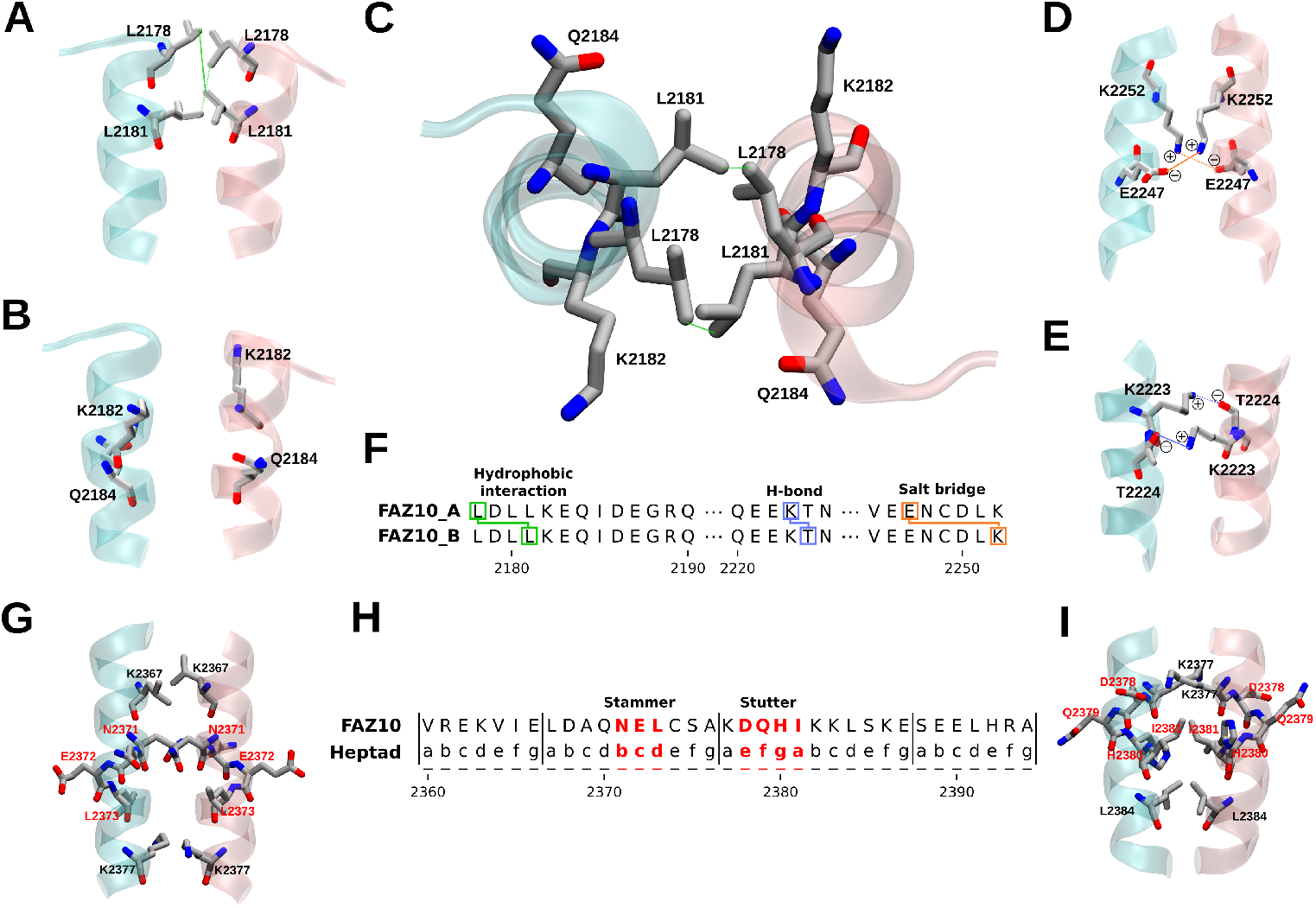
Intermolecular contacts and local heptad disruptions define the architecture of the coiled-coil dimer in the FAZ10 central region. (A) Hydrophobic contacts (*e*.*g*. L2178 and L2181) stabilizing the dimer interface. (B) Residues at the ‘e’ and ‘g’ heptad positions oriented toward the solvent. (C) Close-up view of hydrophobic packing at the dimer interface. (E) Hydrogen bonds involving K2223 and T2224. (F) Sequence schematic highlighting residues contributing to intermolecular interactions.(G) Stammer insertion (N2371–L2373; ‘bcd’) disrupting the canonical heptad repeat. (H) Simplified sequence map indicating the positions of stammer and stutter insertions. (I) Stutter insertion (D2378– I2381; ‘efga’).

Moreover, manual inspection of the structure revealed two localized deviations from the canonical repeat pattern (Figs. 3G–I). Specifically, a three-residue insertion consistent with a stammer N2371, E2372, and L2373 (Fig. 3G), and a four-residue insertion corresponding to a stutter D2378, Q2379, H2380, and I2381 (Fig. 3I), were identified within the coiled-coil domain. These irregularities introduce subtle perturbations in helical alignment that may contribute to the localized bending and axial flexibility observed in the last frame. Therefore, to further investigate whether such structural deviations influence the long-timescale mechanical behavior of the assembly, we extended our analysis using CG-MD simulations under GōMartini 3 approach [18–20]. This approach enabled improved conformational sampling over longer timescales, allowing us to assess the persistence of bending, axial fluctuations, and overall conformational heterogeneity beyond the temporal limits accessible with AA-MD simulations.

### Large-scale MD simulations of the FAZ10 central region

To examine the stability and mechanical behavior of the FAZ10 central region beyond atomistic timescales, we performed five independent 10 *µ*s of CG-MD simulations. The structural parameters remained stable during the MD simulations. For instance, the RMSD rapidly converged and stabilized between ≈2 nm across all replicas (Fig. 4A), indicating preservation of the overall dimeric form over extended timescales. Residue-level fluctuations (RMSF) revealed also localized flexibility, higher in the globular domains and less significant in the coiled-coil segment (Fig. 4B). The radius of gyration (Rg) fluctuated narrowly around ≈17–18 nm (Fig. 4C), while the end-to-end distance (*R*_*ee*_) showed values between ≈42-48 nm (Fig. 4D), indicating axial breathing without large-scale compaction or unfolding. Furthermore, we determined an inter-vector bending angle (*θ*, see definition in Method section) fluctuating around 160° (Figs. 4E and H), which supports the presence of controlled filament bending rather than rigid-rod behavior.

**Fig. 4:**
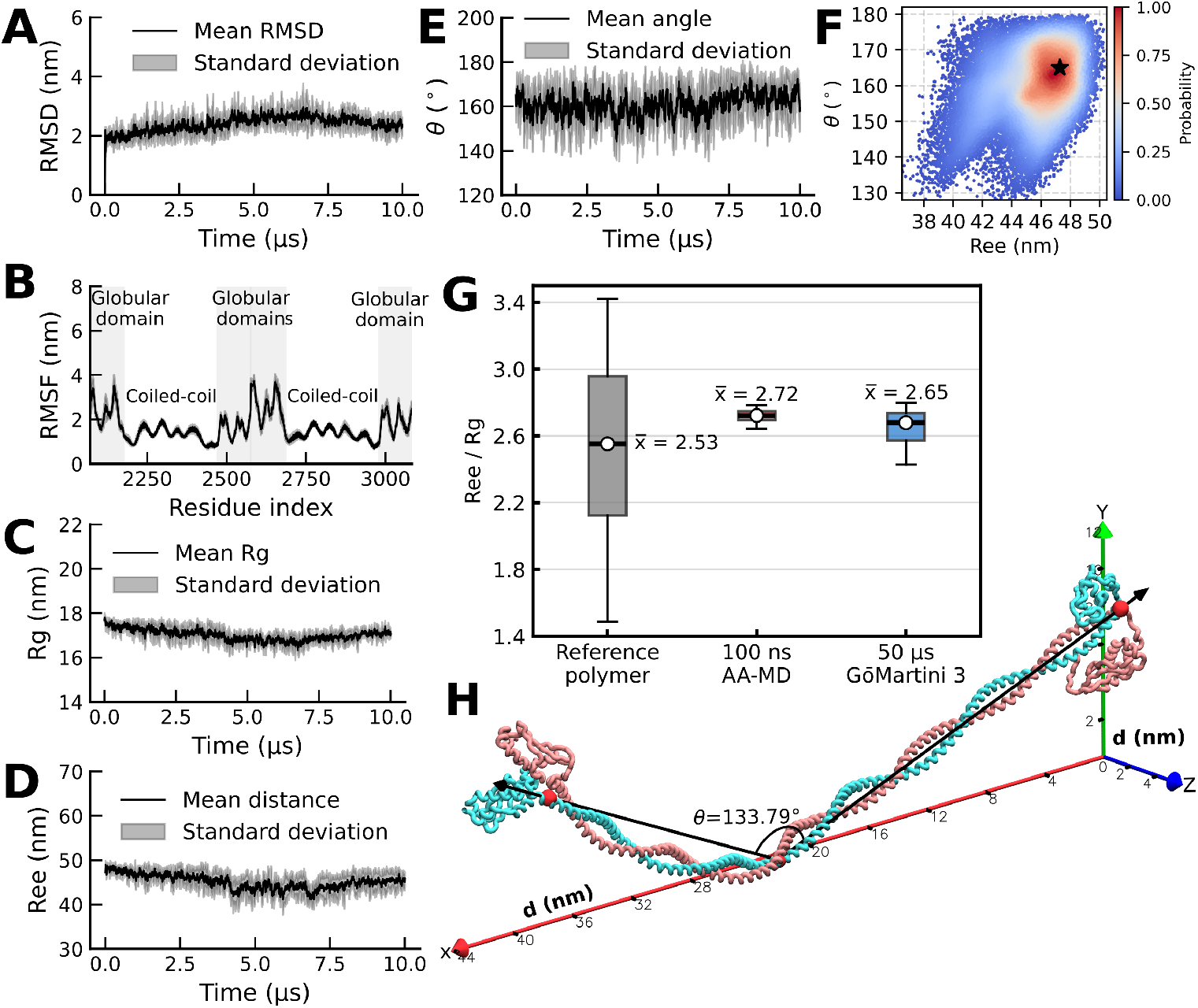
FAZ10 central region exhibits a semi-flexible dimeric architecture with localized bending revealed by large-scale molecular dynamics. (A–D) Mean *±* SD of RMSD, RMSF, *R*_*g*_, and *R*_*ee*_ over time. (E) Time evolution of the bending angle *θ*. (F) Free energy landscape of the FAZ10 central region, based on *P* (*R*_*ee*_, *θ*). The most likely state is highlighted by a black star. (G) Comparison of the *R*_*ee*_*/R*_*g*_ ratio for a reference self-avoiding polymer. (H) Most compact conformation highlighting a localized bend (*θ* = 133.79^*°*^).

The free energy landscape, constructed using *R*_ee_ and *θ*, revealed a single dominant conformational basin centered at approximately *R*_ee_ ≈47 nm and *θ* ≈165° (Fig. 4F), corresponding to the highest-probability region. This probability indicates that the system predominantly assumes an elongated conformation with mild bending, demonstrating that the FAZ10 central region behaves as an elastically deformable filament.

To evaluate whether FAZ10 central region exhibits polymer-like dynamics, we calculated the *R*_ee_/*R*_g_ ratio. This region exhibited mean ratio values of 2.72 in AA-MD simulations and 2.65 in CG-MD simulations, compared to 2.53 for the reference self-avoiding polymer single chain (Fig. 4G). In all cases, the ratio fluctuates, indicating ongoing conformational variability. However, the FAZ10 dimer samples a narrower range of *R*_ee_/*R*_g_ values than expected for a single self-avoiding chain. This suggests that, although the overall dimensions remain within a polymer-like regime, the accessible conformational space is more restricted, likely due to the geometric constraints imposed by dimerization. Furthermore, to illustrate how shortening is accommodated within the CG ensemble, we selected the most contracted conformation (Fig. 4H). In this state, the reduction in *R*_ee_ is achieved primarily through a single localized bend, with a bending angle of *θ* = 133.79^*°*^. This behavior is consistent with a semi-flexible filament in which reductions in *R*_ee_ are accommodated by localized bending rather than global collapse.

### Experimental validation of dimerization of the FAZ10 central region

In this section, we focused on the experimental validation of the FAZ10 central region to corroborate our *in silico* findings. For that, we obtained recombinant proteins corresponding to the central region, the isolated coiled-coil domain, and the isolated globular domain. The purified recombinant proteins were analysed by Size Exclusion Chromatography (SEC) (Fig. 5A). The analytical profile of the central FAZ10 region revealed two major elution peaks, one eluting at the column void volume, indicative of high-molecular-weight aggregates, and a second peak (1) was selected for further analysis (Fig. 5A). Afterward, SDS–PAGE confirmed the expected molecular mass (≈ 66 kDa) for this peak (Fig 5B).

**Fig. 5:**
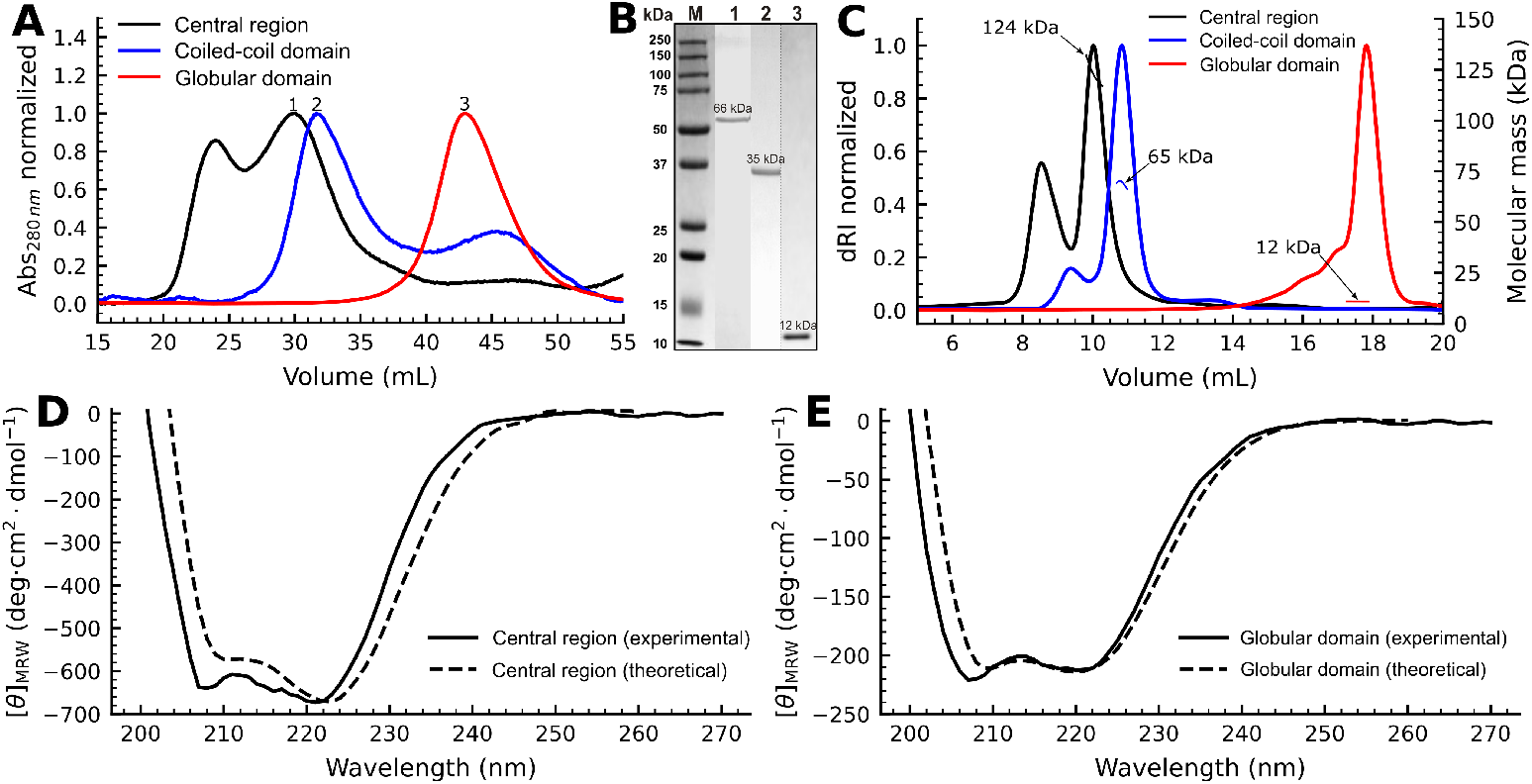
Experimental characterization of recombinant proteins corresponding to the FAZ10 central region. (A) Analytical SEC profiles of the recombinant central region (black), isolated coiled-coil domain (blue), and globular domain (red), monitored at 280 nm. (B) SDS–PAGE of purified fractions stained with Coomassie Blue. M = molecular weight marker; lane 1, central region (66 kDa); lane 2, coiled-coil domain (35 kDa); lane 3, globular domain (12 kDa). (C) SEC–MALS analysis indicates that the central region forms a dimer (124 kDa) with minor higher-order species, the coiled-coil domain is dimeric (65 kDa), and the globular domain is monomeric (12 kDa). (D) Experimental (solid line) and theoretical (dashed line) CD spectra of the central region. The 222/208 ellipticity ratio was 1.04, consistent with a coiled-coil–type *α*-helical arrangement; minima in mean residue ellipticity (MRW) closely coincide between experimental and theoretical spectra. (E) Experimental (solid line) and theoretical (dashed line) CD spectra of the globular domain. The 222/208 ellipticity ratio was 0.95, indicative of a predominantly *α*-helical globular fold; MRW minima similarly overlap between experimental and theoretical spectra.

Similarly, SEC profiles of the coiled-coil domain also displayed two peaks, with the dominant fraction corresponding to the expected molecular weight (≈35 kDa), whereas the globular domain exhibited a single peak corresponding to a ≈12 kDa protein. However, SEC alone did not allow for an unambiguous determination of oligomeric state. For this reason, to accurately assess molecular mass, SEC coupled with multi-angle light scattering (SEC–MALS) was performed (Fig. 5C). The central region displayed a predominant peak with an estimated molecular mass of ≈124 kDa, approximately twice its monomeric mass, which is compatible with dimer formation. The coiled-coil domain also exhibited a molecular mass of ≈65 kDa, providing further evidence for dimerization, while the globular domain remained monomeric (≈12 kDa).

To assess the secondary structure of the central region and the globular domain, we applied Circular Dichroism (CD) spectroscopy and compared experimental data with theoretical spectra (Fig. 5D-E). Both spectra of the central region exhibited characteristic minima at ≈208 nm and 222 nm, with a slightly deeper minimum at ≈222 nm, indicative of a coiled-coil–rich *α*-helical architecture. The ≈222/208 nm ellipticity ratio of 1.04 further supports a coiled-coil–dominated organization [21] (Fig. 5D). The globular domain also exhibited minima at 208 nm and 222 nm, but with a more pronounced minimum at 208 nm and a lower 222/208 ratio (0.95), consistent with a compact mixed *α*/*β* fold, different from the spectroscopic signal of coiled-coil assemblies. Finally, the theoretical curves presented good agreement for both the central region and its globular domain constructs.

## Discussion

This study presents the structural organization of the FAZ10 central region of *T. brucei*. This region displays lower disorder and high sequence conservation across *T. brucei* strains, suggesting that the structural features described here are likely preserved across variants. The model reveals an elongated architecture composed of a parallel coiled-coil dimer flanked by globular domains. The alignment of residues at the ‘a’ and ‘d’ positions along the interface, together with classic knobs-into-holes packing, indicates that this region follows the canonical principles of coiled-coil organization [22, 23]. In addition, residues at the ‘e’ and ‘g’ positions form a peripheral electrostatic network characteristic of parallel coiled-coils [24, 25]. Together, these features provide structural evidence that the central region assembles as a canonical coiled-coil homodimer rather than a nonspecific helical association.

Biophysical analyses further support this interpretation. Both the full central region and the isolated coiled-coil domain predominantly assemble as stable dimers in solution, while a reproducible minor population elutes at the void volume in SEC and SEC–MALS. This behavior is consistent with the intrinsic self-association tendencies of extended coiled-coil interfaces [24]. The recurrence of this higher-order species across independent preparations indicates a reproducible tendency for self-association beyond the dimeric state. Such behavior is often observed in coiled-coil proteins that form higher-order assemblies associated with filamentous scaffolds.

The identification of stammers and stutters indicates that the coiled-coil is architecturally tuned rather than uniformly rigid. These heptad discontinuities are recognized modulators of local helical geometry, introducing controlled deviations that allow bending or torsional adjustment within otherwise stable assemblies [26–28]. In the FAZ10 protein, the spatial proximity of a stammer and a compensatory stutter likely maintains overall helical continuity while locally adjusting superhelical parameters [28]. This balance between structural persistence and conformational adaptability is likely essential for FAZ proteins, which must undergo mechanical forces generated during flagellar beating, cell elongation, and cytokinesis [9, 13].

Thus, the FAZ10 central region appears optimized to combine mechanical resilience with controlled flexibility within the dynamic FAZ cytoskeleton. Beyond flexibility, the parallel dimeric configuration may directly contribute to mechanical tension within the FAZ filament, as parallel coiled-coils behave as semi-flexible elastic elements capable of sustaining and distributing longitudinal forces while permitting limited extension under load.

In this context, FAZ10 may function as a load-bearing fiber within the FAZ staples, where its parallel architecture enables a coordinated mechanical response along the longitudinal axis of the filament and across the flagellum–cell body connection. This semi-flexibility may also facilitate interactions with FAZ-associated partners; notably, FAZ10 has been proposed to interact with components such as ClpGM6 [14]. Such adaptability could further support coupling between the paraflagellar rod and the FAZ filament [14]. The dimeric arrangement, reinforced by interactions at both the core and periphery, provides a structural basis for coupling elasticity with stability, supporting a model in which FAZ10 contributes actively to force transmission rather than acting as a passive scaffold.

Analysis by SEC–MALS further delineates the structural hierarchy for the central region. Dimerization is mediated primarily by the coiled-coil domain, whereas the isolated globular domains remain monomeric in solution. Although AlphaFold2 and MD simulations predict spatial proximity between globular segments, such contacts are insufficient to drive stable association under the conditions tested. Instead, oligomerization arises from the parallel association of the extended helices, consistent with canonical coiled-coil architecture [22]. The globular domains, while not contributing to dimerization, may serve alternative roles within the full-length protein. In the absence of defined biochemical functions, it is plausible that these regions participate in mediating interactions between FAZ10 dimers or between FAZ10 and other FAZ components. Such interaction interfaces remain to be experimentally characterized.

Given that the central segment is embedded within the full-length FAZ10 protein, it is reasonable to propose that dimerization extends to the full-length molecule, primarily mediated by its coiled-coil regions. Previous analyses identified additional predicted coiled-coil segments throughout FAZ10 [14], consistent with elongated densities observed in filamentous FAZ components [29]. These observations support a model in which FAZ10 contributes to the longitudinal architecture of the FAZ cytoskeleton through an extended, coiled-coil–rich framework that supports higher-order organization.

More broadly, coiled-coil–mediated dimerization represents a recurring feature among the FAZ network, including CC2D [30], TbBILBO1 [31], and CIF3 [12]. The convergence of similar structural modules across multiple FAZ components suggests a shared assembly in which elongated dimers function as modular elements within an interconnected cytoskeletal network. In this structural framework, FAZ10 may act as a longitudinal organizer that integrates elasticity, tension transmission, and spatial coordination during cell morphogenesis.

Together, these findings establish the first structural basis for the FAZ10 protein and define principles by which extended parallel coiled-coil dimers contribute to the mechanical and architectural properties of the FAZ cytoskeleton in *T. brucei*.

## Methods

### Cloning, expression, and purification

Primers (Table S1) were designed to amplify the central region of FAZ10 (Tb427.07.3330) from *Trypanosoma brucei* strain 427. The full central region, as well as the isolated coiled-coil and globular domains, were PCR-amplified and cloned into pET-28a-TEV or pET-28a-SUMO vectors, following appropriate double digestion (BamHI/XhoI for SUMO; NdeI/XhoI for TEV). Recombinant proteins were expressed in *E. coli* BL21 Rosetta™(DE3) cells cultured in Luria–Bertani (LB) broth supplemented with chloramphenicol (34 *µ*g·mL^*−*1^) and kanamycin (50 *µ*g·mL^*−*1^). When cultures reached an OD_600_ of 0.6, protein expression was induced with isopropyl 1-thio-*β*-D-galactopyranoside (IPTG) at a final concentration of 0.1 mM for the central region domains or 1 mM for the full central region. Cultures were incubated for 4 h at 37 ^*°*^C with shaking. Subsequently, cells were collected by centrifugation (10,000 *g* for 30 min, at 4 ^*°*^C), and suspended in ice-cold lysis buffer (50 mM Tris–HCl pH 7.4, 300 mM NaCl, 20 mM imidazole, 1 mg/mL lysozyme, 0.05% Tween-20, containing 1 mM DTT, 1 mM benzamidine, 0.01 mM leupeptin, and 1 mM PMSF). Cell lysis was performed by sonication using a QSonica Q125 Sonicator (M2 Scientifics) with 30 s on / 30 s off cycles for a total of eight pulses and centrifuged at 10,000 RPM for 30 min at 4 ^*°*^C.

The supernatant was collected and loaded onto an equilibrated Ni–NTA agarose column (GE Healthcare). The column was washed with 10 volumes of lysis buffer, and the target proteins were eluted with lysis buffer containing 100 mM and 250 mM imidazole. SUMO fusion tags were removed from recombinant proteins (coiled-coil and globular domains) by cleavage with SUMO protease (Thermo Fisher Scientific) according to the manufacturer’s instructions [32]. Purification fractions were subsequently analyzed by SDS–PAGE.

### Size Exclusion Chromatography (SEC)

Size Exclusion Chromatography (SEC) was performed using ÄKTA Purifier 10 system (GE Healthcare Life Sciences) coupled to a Superdex 200 column. Fractions were collected at a flow rate of 1 mL/min, with absorbance monitored at 280 nm. Prior to purification. The column was equilibrated with lysis buffer (50 mM Tris–HCl (pH 7.4) and 150 mM NaCl), and data were plotted on a graph using Origin 2024 software (version 10.1.0.178). Fractions were concentrated using Amicon^®^ Ultra centrifugal filter device (mol. wt. cut-off, 3 and 30 kDa, Merck Millipore, Darmstadt, Germany), analyzed in 15% SDS–PAGE, and gels were stained with Coomassie blue. Gel images were acquired using a ChemiDoc XRS+ System (Bio-Rad) and processed with ImageJ.

### Circular Dichroism spectroscopy

Recombinant FAZ10 proteins were monitored by circular dichroism (CD) spectroscopy using a J–815 Jasco spectropolarimeter at the São Carlos Institute of Physics (IFSC-USP). Thermal denaturation was measured using 0.1 cm quartz cuvettes by monitoring ellipticity changes at 222 nm while heating samples (3 *µ*M in 20 mM sodium phosphate pH 7.8, 100 mM NaCl, 4% glycerol) from 10 to 90 °C at a rate of 10 °C/h. The fraction of denatured protein (*f*_*d*_) was calculated as:

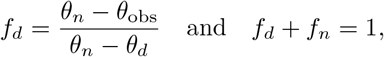

in which *θ*_obs_ is the ellipticity at 222 nm obtained at a particular temperature, *θ*_*d*_ and *θ*_*n*_ are the ellipticities of the denatured and native states, respectively, and *f*_*n*_ is the fraction of protein in the native state. Far-UV CD spectra were processed using CDTool [33] and normalized to mean residue ellipticity based on protein concentration and residue number. Theoretical CD spectra were calculated from AlphaFold2 [17] models using the PDBMD2CD web server [34] and scaled to experimental spectra at 222 nm for comparison. The *θ*222/*θ*208 ratio was calculated from baseline-corrected mean residue ellipticity values at 222 and 208 nm.

### Size-Exclusion Chromatography coupled with Multi-Angle Light Scattering assay (SEC–MALS)

The oligomeric state was characterized using SEC coupled with a miniDAWN^®^ TREOS^®^ multi-angle light scattering detector and an Optilab T–rEX differential refractometer (Wyatt Technology). Aliquots of 20 *µ*M protein were loaded onto a Superdex 200 5/150 column (Cytiva) and analyzed using a high-performance liquid chromatography (HPLC) system (Waters, Milford, MA, USA) consisting of a pump and controller (Waters 600). Chromatography was conducted at a continuous flow rate of 0.3 mL/min in 50 mM HEPES, pH 8.0, and 300 mM NaCl. Coupled with the chromatographic separation, the DAWN TREOS miniature system (Wyatt Technology) was employed to determine mass distribution, size, and composition, while the Opti-Lab T-REX system (Wyatt Technology) measured the differential refractive index. Data were processed using ASTRA7 software (Wyatt Technology), and the graphs were generated and plotted in Origin 2024 (v. 10.1.0.178).

### Sequence analysis

The FAZ10 sequence (Tb927.7.3330) was obtained from TriTrypDB [35]. Secondary structure and intrinsic disorder were analyzed using PSIPRED [36] and IUPred2A [37](long disorder mode), respectively. Coiled-coil domains, heptad repeats, and oligomeric states were predicted using the MARCOIL algorithm [16] via the Waggawagga server [38].

### Protein modeling

The central region was modeled *de novo* using ColabFold v1.5.2 (AlphaFold2) [17, 39]. MMseqs2 (Uniref+environmental) msa mode, automatic model type, unpaired+paired pair mode, and auto run recycles were selected as the input parameters, obtaining five final models, and the best-ranked model was relaxed using AMBER force field by Google Colab [39]. The quality of the model was evaluated using pLDDT and PAE metrics.

### All-atom molecular dynamics simulations

All-atom MD simulations of the central region of the FAZ10 protein (507 aa) were performed in GROMACS v. 2023.5 [40] using the Amber ff14SB force field [41]. The protein was centered and aligned along its principal axes in a rectangular simulation box (55 ×10 × 10 nm^3^). The system was composed of 16,194 atoms. Then, the system was solvated with TIP3P water and ionized with Na^+^/Cl^*−*^ to neutralize the system and reach a physiological ionic concentration of 0.15 M. The solvated and ionized system was composed of 537,764 atoms. Following, the system was energy-minimized using the steepest-descent algorithm for up to 50,000 steps, with a minimization step size of 0.01 nm and a convergence criterion of 1000 kJ/mol/nm. Temperature was equilibrated at 300 K using the velocity-rescale thermostat [42] over 100 ps. Pressure was equilibrated at 1 bar using the Parrinello-Rahman barostat [43] over 100 ps. During minimization and equilibration steps, harmonic restraints of 10 kcal/mol/Å were applied to C-alpha atoms. The LINCS algorithm [44] was used to constrain all bonds, and long-range electrostatics were treated with the particle-mesh Ewald (PME) method [45]. Production was performed for a time of 100 ns with 3 replicas.

### Coarse-grained molecular dynamics simulations

Coarse-grained MD simulations [46] are advanced tools to overcome time and length scale limitations present in AA-MD. Here we employed the Martini 3 [47] force field for proteins and the GōMartini 3 approach to retain secondary and tertiary structure [48]. To enhance conformational space exploration of FAZ10, the contact map determination criteria [49, 50] and the corresponding optimization protocol [51] were employed to capture the protein flexibility in Martini 3 simulations. MD simulations employed GROMACS v.2023.5. The All-atom MD trajectories were used to build the optimized contact map (Fig. S6). The frame with the maximum contact count over the trajectories was selected as the initial conformation for subsequent GōMartini 3 simulations. Here, the protein assembly was centered in a rectangular box (55 × 15 × 15 nm^3^). The CG system was composed of 2258 CG beads. The system topology was built using the *martinize2* algorithm [52]. The protein was subjected to vacuum energy minimization for 5000 steps using the steepest descent algorithm [53]. Then, the system was solvated using Martini 3 force field. Following, the system was neutralized by adding Na^+^/Cl^*−*^ ions to reach a concentration of 0.15 M. The final solvated and ionized system was composed of 103,457 CG beads. A second energy minimization step was performed for the solvated and neutralized system for 5000 steps. During equilibration steps, we employed an integration time step of 20 fs, and position restraints were applied to backbone beads (BB) in the protein. Equilibration was conducted in two stages. The first in the NVT ensemble at 300 K using the V-rescale thermostat [42], with a coupling time of 1.0 ps. This phase lasted 5 ns. The second equilibration step was carried out in the NPT ensemble at 1 atm using the C-rescale thermostat [54], with isotropic pressure coupling equal to 10^*−*4^ bar^*−*1^ and a pressure coupling time constant of 12 ps during 10 ns. Production was conducted under NPT conditions. Five replicas of 10 *µ*s each (totaling 50 *µ*s) were produced.

### Self-avoiding polymer reference simulations

To obtain a reference distribution for the *R*_*ee*_*/R*_*g*_ ratio (see Structural analysis section), we performed MD dynamics simulations using the ESPResSo package [55] to model a self-avoiding single-chain polymer. The system comprised a chain of 507 beads in a cubic periodic box, with the box length determined from a low monomer number density (*ρ* = 0.001) to minimize finite-size effects. Consecutive beads were connected by harmonic bonds (*k* = 100, *r*_0_ = 1.0, reduced units), while excluded-volume interactions were described by a purely repulsive Weeks–Chandler–Andersen (WCA) potential with *σ* = 1.5 and a cutoff at 2^1*/*6^*σ*. Initial polymer conformations were generated as self-avoiding walks (SAW) enforcing fixed bond lengths and a minimum separation between non-neighboring beads. The equations of motion were integrated using a velocity-Verlet scheme under Langevin dynamics at temperature *T* = 1.0 and friction coefficient *γ* = 1.0. To remove unfavorable initial contacts, simulations were preceded by an equilibration phase using a reduced time step, increased damping, force capping, and with excluded-volume interactions temporarily disabled, followed by a gradual ramping of the WCA interaction strength to its target value. Production simulations were performed for 2 ×10^6^ integration steps with a time step of 0.02. Polymer conformational observables, including the *R*_*ee*_ distance, *R*_g_, and their ratio, were computed at regular intervals from particle coordinates. Trajectories were stored in VTF format for visualization, and reduced simulation units were mapped to physical units using *σ* = 0.47 nm and a time conversion factor of 0.25 ns per reduced time unit.

### Structural analysis

Molecular interactions were identified via the PLIP algorithm [56]. System stability was evaluated using the root-mean-square deviation (RMSD), root-mean-square fluctuation (RMSF), radius of gyration (*R*_*g*_), and end-to-end distance (*R*_*ee*_). The *R*_*g*_, was calculated with GROMACS as the mass-weighted root-mean-square distance of the BB atoms of the system from their center of mass:

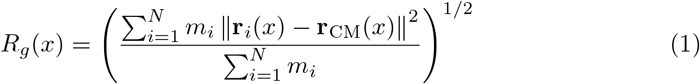

where *N* is the number of selected BB atoms, *m*_*i*_ is the mass of particle *i*, **r**_*i*_(*x*) is the position vector of particle *i* in configuration *x*, and **r**_CM_(*x*) is the center-of-mass position vector of the selected group. While the *R*_*ee*_ was calculated using a TCL script that solves the following equation:

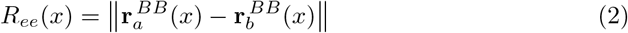

where 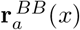 and 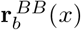 are the position vectors of the selected terminal BB atoms (resid Asp2070 and Pro2576) in configuration *x*.

As a collective variable, we defined the inter-vector angle fluctuation (*θ*) between two vectors: **v**_1_, constructed from the BB atoms of residues Asp2179 and Glu2340, and **v**_2_, constructed from the BB atoms of residues Glu2340 and Asn2465 (see Figure 4H). The angle between the vectors was solved with the dot-product definition:

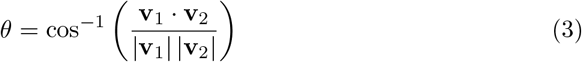

where **v**_1_ · **v**_2_ is the scalar product between the two vectors, and |**v**_1_| and |**v**_2_| are their magnitudes. This definition provides the instantaneous angle between the two segments formed by Asp2179–Glu2340 and Glu2340–Asn2465, thereby quantifying local bending motions in this region. Lower *θ* values indicate a more aligned or extended arrangement of the two vectors, whereas larger *θ* values reflect a more pronounced bend. This calculation was performed by a TCL script in VMD.

The probability landscape (or FEL) of the system was then computed using *R*_*ee*_ and *θ* as collective variables. To support the model-derived conformational trends, we further evaluated the *R*_*ee*_*/R*_*g*_ relationship as a reference metric of polymer-like behavior (see MD simulation methods). For an ideal Gaussian polymer chain, the *R*_*ee*_ and *R*_*g*_ are related by:

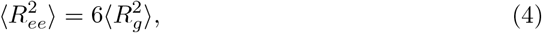

which implies

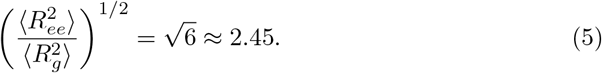

Accordingly, we computed the *R*_*ee*_*/R*_*g*_ distributions across all independent replicas for both AA-MD and GōMartini 3 simulations.

Finally, following FAIR principles in biomolecular simulations [57], the AA and CG trajectories, together with the raw data derived from the MD simulations and experimental analyses, are publicly available through the Zenodo repository: https://doi.org/10.5281/zenodo.19234680.

## Acknowledgements

The authors acknowledge Prof. Dr. Richard Garratt (Structural Biology Laboratory, Institute of Physics of São Carlos, University of São Paulo) for access to laboratory facilities and valuable suggestions during manuscript preparation, and Andressa Alves Pinto for her assistance with SEC–MALS measurements. We also thank Dr. Luis F. Cofas-Vargas for early discussions on the MD simulations. A.B.P. acknowledges financial support from the National Science Center, Poland, under grant 2022/45/B/NZ1/02519, and gratefully acknowledges the Polish high-performance computing infrastructure PLGrid (HPC Center: ACK Cyfronet AGH) for providing computer facilities and support within computational grant no. PLG/2024/017332 and PLG/2025/018510. D.A.L. acknowledges the São Paulo Research Foundation (FAPESP; 2021/08158-9). This work was supported by the São Paulo Research Foundation (FAPESP; 2018/03677-5 and 2023/17920-7) to MMAB, by Fundação de Apoio ao Ensino, Pesquisa e Assistência (FAEPA) to MMAB, and by the Coordination for the Improvement of Higher Education Personnel (CAPES; grant no. 88887.816952/2023-00) to C.M.O.-M.

## Author contributions

C.M.O.-M.: Conceptualization, Methodology, Investigation, Data curation, Formal analysis, Validation, Visualization, Writing – original draft, Writing – review & editing.

D.A.L.: Investigation, Data curation, Formal analysis, Writing – original draft, Writing – review & editing, Supervision. C.I.: Investigation, Formal analysis, Validation, Writing – review & editing. L.C.A.: Investigation, Formal analysis, Validation. G.E.O.-R.: Investigation, Data curation, Formal analysis, Software, Visualization, Writing – original draft, Writing – review & editing. A.B.P.: Methodology, Investigation, Writing – review & editing, Supervision, Funding acquisition. M.M.A.B.: Conceptualization, Methodology, Formal analysis, Writing – original draft, Writing – review & editing, Supervision, Funding acquisition. All authors read and approved the final manuscript.

## Source data

Source data in Excel format available.

## References

[1] The Lancet Microbe. Neglected tropical diseases: golden age of elimination? The Lancet Microbe (2025).

[2] World Health Organization. Ending the neglect to attain the Sustainable Development Goals: A road map for neglected tropical diseases 2021–2030 (World Health Organization, Geneva, Switzerland, 2021). URL https://www.who.int/publications/i/item/9789240010352.

[3] Hotez, P. J. et al. Control of neglected tropical diseases. New England Journal of Medicine 357, 1018–1027 (2007).

[4] Büscher, P., Cecchi, G., Jamonneau, V. & Priotto, G. Human african trypanosomiasis. The Lancet 390, 2397–2409 (2017).

[5] Kohl, L. & Gull, K. Molecular architecture of the trypanosome cytoskeleton. Molecular and Biochemical Parasitology 93, 1–9 (1998).

[6] Matthews, K. R. The developmental cell biology of trypanosoma brucei. Journal of Cell Science 118, 283–290 (2005).

[7] Lee, K. J., Zhou, Q. & Li, Z. Crk2 controls cytoskeleton morphogenesis in Trypanosoma brucei by phosphorylating β-tubulin to regulate microtubule dynamics. PLoS Pathog. 19, e1011270 (2023).

[8] Kohl, L., Sherwin, T. & Gull, K. Assembly of the paraflagellar rod and the flagellum attachment zone complex during the trypanosoma brucei cell cycle. Journal of eukaryotic microbiology 46, 105–109 (1999).

[9] Vaughan, S., Kohl, L., Ngai, I., Wheeler, R. J. & Gull, K. A repetitive protein essential for the flagellum attachment zone filament structure and function in trypanosoma brucei. Protist 159, 127–136 (2008).

[10] Sunter, J. D., Varga, V., Dean, S. & Gull, K. A dynamic coordination of flagellum and cytoplasmic cytoskeleton assembly specifies cell morphogenesis in trypanosomes. Journal of Cell Science 128, 1580–1594 (2015).

[11] Kohl, L., Robinson, D. & Bastin, P. Novel roles for the flagellum in cell morphogenesis and cytokinesis of trypanosomes. The EMBO Journal 22, 5336–5346 (2003).

[12] Zhou, Q., Hu, H., He, C. Y. & Li, Z. Assembly and maintenance of the flagellum attachment zone filament in trypanosoma brucei. Journal of Cell Science 128, 2361–2372 (2015).

[13] Sunter, J. D. & Gull, K. The flagellum attachment zone: The Cellular Ruler of trypanosome morphology. Trends in Parasitology 32, 309–324 (2016).

[14] Moreira, B. P., Fonseca, C. K., Hammarton, T. C. & Baqui, M. M. A. Giant faz10 is required for flagellum attachment zone stabilization and furrow positioning in Trypanosoma brucei. Journal of Cell Science 130, 1179–1193 (2017).

[15] McGuffin, L. J., Bryson, K. & Jones, D. T. The psipred protein structure prediction server. Bioinformatics 16, 404–405 (2000).

[16] Delorenzi, M. & Speed, T. An hmm model for coiled-coil domains and a comparison with pssm-based predictions. Bioinformatics 18, 617–625 (2002).

[17] Jumper, J. et al. Highly accurate protein structure prediction with alphafold. Nature 596, 583–589 (2021).

[18] Poma, A. B., Cieplak, M. & Theodorakis, P. E. Combining the Martini and structure-based coarse-grained approaches for molecular dynamics studies of conformational transitions in proteins. J. Chem. Theory Comput. 13, 1366–1374 (2017).

[19] Souza, P. C. T. et al. GōMartini 3: From large conformational changes in proteins to environmental bias corrections. Nat. Commun. 16, 4051 (2025).

[20] Cofas-Vargas, L. F., Moreira, R. A., Poblete, S., Chwastyk, M. & Poma, A. B. The go-martini approach: Revisiting the concept of contact maps and the modelling of protein complexes. Acta Phys. Pol. A. 145, S9 (2024).

[21] Sala, F. A., Valadares, N. F., Macedo, J. N. A., Borges, J. C. & Garratt, R. C. Heterotypic coiled-coil formation is essential for the correct assembly of the septin heterofilament. Biophysical Journal 111, 2608–2619 (2016).

[22] Truebestein, L. & Leonard, T. A. Coiled-coils: The long and short of it. BioEssays 38, 903–916 (2016).

[23] Lupas, A. N. & Bassler, J. Coiled coils: A model system for the 21st century. Trends in Biochemical Sciences 42, 130–140 (2017).

[24] Lupas, A. N. & Gruber, M. in The structure of-helical coiled coils, Vol. 70 of Advances in Protein Chemistry 37–38 (Academic Press, 2005).

[25] Woolfson, D. N. Coiled-Coil Design: Updated and Upgraded, 35–61 (Springer International Publishing, Cham, 2017).

[26] Brown, J. H., Cohen, C. & Parry, D. A. D. Heptad breaks in α-helical coiled coils: Stutters and stammers. Proteins: Structure, Function, and Bioinformatics 26, 134–145 (1996).

[27] Gruber, M. & Lupas, A. N. Historical review: Another 50th anniversary – new periodicities in coiled coils. Trends in Biochemical Sciences 28, 679–685 (2003).

[28] Schmidt, N. W., Grigoryan, G. & DeGrado, W. F. The accommodation index measures the perturbation associated with insertions and deletions in coiled-coils: Application to understand signaling in histidine kinases. Protein Science 26, 414–435 (2017).

[29] Trépout, S. In situ structural analysis of the flagellum attachment zone in trypanosoma brucei using cryo-scanning transmission electron tomography. Journal of Structural Biology: X 4, 100033 (2020).

[30] Zhou, Q., Liu, B., Sun, Y. & He, C. Y. A coiled-coil- and c2-domain-containing protein is required for faz assembly and cell morphology in trypanosoma brucei. Journal of Cell Science 124, 3848–3858 (2011).

[31] Florimond, C. et al. Bilbo1 is a scaffold protein of the flagellar pocket collar in the pathogen trypanosoma brucei. PLoS pathogens 11, e1004654 (2015).

[32] Invitrogen. Champion™ pET SUMO Protein Expression System User Manual. Carlsbad, CA, USA (2010). Rev. Date: 18 June 2010.

[33] Lees, J., Smith, B., Wien, F., Miles, A. & Wallace, B. Cdtool—an integrated software package for circular dichroism spectroscopic data processing, analysis, and archiving. Analytical Biochemistry 332, 285–289 (2004).

[34] Drew, E. D. & Janes, R. W. Pdbmd2cd: providing predicted protein circular dichroism spectra from multiple molecular dynamics-generated protein structures. Nucleic Acids Research 48, W17–W24 (2020).

[35] Amos, B. et al. VEuPathDB: the eukaryotic pathogen, vector and host bioinformatics resource center. Nucleic Acids Res. 50, D898–D911 (2021).

[36] Jones, D. T. Protein secondary structure prediction based on position-specific scoring matrices. J. Mol. Biol. 292, 195–202 (1999).

[37] Mészáros, B., Erdős, G. & Dosztányi, Z. Iupred2a: context-dependent prediction of protein disorder as a function of redox state and protein binding. Nucleic Acids Res. 46, W329–W337 (2018).

[38] Simm, D., Hatje, K. & Kollmar, M. Waggawagga: comparative visualization of coiled-coil predictions and detection of stable single α-helices (sah domains). Bioinformatics 31, 767–769 (2014).

[39] Mirdita, M. et al. Colabfold: making protein folding accessible to all. Nat. Methods 19, 679–682 (2022).

[40] Abraham, M. et al. Gromacs: high performance molecular simulations through multi-level parallelism from laptops to supercomputers. SoftwareX 1-2, 19–25 (2015).

[41] Maier, J. et al. Ff14sb: improving the accuracy of protein side chain and backbone parameters from ff99sb. J. Chem. Theory Comput. 11, 3696–3713 (2015).

[42] Bussi, G., Donadio, D. & Parrinello, M. Canonical sampling through velocity rescaling. J. Chem. Phys. 126, 014101 (2007).

[43] Parrinello, M. & Rahman, A. Polymorphic transitions in single crystals: A new molecular dynamics method. J. Appl. Phys. 52, 7182–7190 (1981).

[44] Hess, B., Bekker, H., Berendsen, H. J. C. & Fraaije, J. G. E. M. Lincs: A linear constraint solver for molecular simulations. J. Comput. Chem. 18, 1463–1472 (1997).

[45] Petersen, H. G. Accuracy and efficiency of the particle mesh ewald method. J. Chem. Phys. 103, 3668–3679 (1995).

[46] B. Poma, A. et al. Recent advances in machine learning and coarse-grained potentials for biomolecular simulations and their applications. Biophysical Journal 125, 327–343 (2025).

[47] Souza, P. C. T. et al. Martini 3: a general purpose force field for coarse-grained molecular dynamics. Nat. Methods 18, 382–388 (2021).

[48] Souza, P. C. et al. Gōmartini 3: From large conformational changes in proteins to environmental bias corrections. Nature Communications 16, 4051 (2025).

[49] Poma, A. B., Thu, T. T. M., Tri, L. T. M., Nguyen, H. L. & Li, M. S. Nanomechanical stability of aβ tetramers and fibril-like structures: Molecular dynamics simulations. The Journal of Physical Chemistry B 125, 7628–7637 (2021).

[50] Cofas-Vargas, L. F., Olivos-Ramirez, G. E., Marrink, S. J. & Poma, A. B. A comparative nanomechanical study of antibody and nanobody binding to sars-cov-2 variants. Physical Chemistry Chemical Physics (2026).

[51] Olivos-Ramirez, G. E., Cofas-Vargas, L. F., Marrink, S. J. & Poma, A. B. An optimized contact map for GōMartini 3 enabling conformational changes in protein assemblies. bioRxiv (2025). Preprint.

[52] Kroon, P. C. et al. Martinize2 and vermouth: Unified framework for topology generation. eLife (2025).

[53] Wardi, Y. A stochastic steepest-descent algorithm. Journal of Optimization Theory and Applications 59, 307–323 (1988).

[54] Bernetti, M. & Bussi, G. Pressure control using stochastic cell rescaling. J. Chem. Phys. 153, 114107 (2020).

[55] Weik, F. et al. Espresso 4.0: an extensible software package for simulating soft matter systems. Eur. Phys. J. Spec. Top. 227, 1789–1816 (2019).

[56] Schake, P., Bolz, S. N., Linnemann, K. & Schroeder, M. PLIP 2025: introducing protein–protein interactions to the protein–ligand interaction profiler. Nucleic Acids Res. 53, W463–W465 (2025).

[57] Amaro, R. E. et al. The need to implement fair principles in biomolecular simulations. Nature methods 22, 641–645 (2025).

