## Supplementary Figures and Tables for "Structural modelling and biophysical analyses reveal a dimeric coiled-coil architecture in the FAZ10 central region of *Trypanosoma brucei*"

Table S1: Primers designed for the cloning and expression of recombinant proteins from the central region of FAZ10.

| **Construct** | **Primer** | **Sequence (5'–3')** |
|---|---|---|
| Globular domain of the central region of FAZ10 | Forward | TATTATCATATGGATTCTCGGTGGATGGCT |
| | Reverse | TATTATCTCGAGTTAAGGTTCATTTAATTTCAACA |
| Coiled-coil domain of the central region of FAZ10 | Forward | TATTATGGATCCAAGGATGGCCTTGATCTATT |
| | Reverse | TATTATCTCGAGTTACCACCGAGAATCATGTTT |
| Central region of FAZ10 | Forward | TATTATGGATCCGATTCTCGGTGGATGGCTTC |
| | Reverse | TATTATCTCGAGTTAAGGTTCATTTAATTTCAACA |

Table S2: Hydrophobic interactions at the FAZ10 central region dimer interface identified by PLIP.

| Chain B | Chain A | Position | Distance (Å) | Chain B | Chain A | Position | Distance (Å) |
|---|---|---|---|---|---|---|---|
| Tyr2159 | Asp2070 | Tyr–Asp | 3.63 | Leu2178 | Leu2178 | Leu–Leu | 3.93 |
| Leu2181 | Lys2182 | Leu–Lys | 3.73 | Leu2181 | Leu2181 | Leu–Leu | 3.98 |
| Leu2181 | Ile2185 | Leu–Ile | 3.50 | Ile2185 | Ile2185 | Ile–Ile | 3.81 |
| Gln2191 | Leu2192 | Gln–Leu | 3.71 | Leu2192 | Leu2192 | Leu–Leu | 3.69 |
| Leu2192 | Leu2195 | Leu–Leu | 3.54 | Leu2195 | Leu2199 | Leu–Leu | 3.67 |
| Leu2199 | Leu2195 | Leu–Leu | 3.75 | Leu2199 | Glu2198 | Leu–Glu | 3.78 |
| Leu2199 | Leu2199 | Leu–Leu | 3.92 | Leu2202 | Leu2199 | Leu–Leu | 3.34 |
| Leu2202 | Leu2202 | Leu–Leu | 3.86 | Leu2202 | Leu2206 | Leu–Leu | 3.98 |
| Val2209 | Leu2206 | Val–Leu | 3.76 | Val2209 | Val2209 | Val–Val | 3.69 |
| Glu2212 | Arg2213 | Glu–Arg | 4.00 | Arg2213 | Val2209 | Arg–Val | 3.62 |
| Leu2216 | Arg2213 | Leu–Arg | 3.72 | Leu2216 | Leu2216 | Leu–Leu | 3.88 |
| Leu2216 | Leu2217 | Leu–Leu | 3.69 | Leu2217 | Leu2216 | Leu–Leu | 3.58 |
| Lys2223 | Lys2223 | Lys–Lys | 3.86 | Leu2227 | Lys2223 | Leu–Lys | 3.79 |
| Leu2227 | Leu2227 | Leu–Leu | 3.75 | Leu2227 | Gln2230 | Leu–Gln | 3.71 |
| Gln2230 | Leu2227 | Gln–Leu | 3.79 | Leu2234 | Val2237 | Leu–Val | 3.90 |
| Val2237 | Val2237 | Val–Val | 3.49 | Val2237 | Leu2241 | Val–Leu | 3.68 |
| Leu2241 | Leu2244 | Leu–Leu | 3.46 | Leu2244 | Leu2241 | Leu–Leu | 3.77 |
| Leu2244 | Leu2244 | Leu–Leu | 3.71 | Leu2244 | Val2245 | Leu–Val | 3.41 |
| Val2245 | Leu2244 | Val–Leu | 3.99 | Asn2248 | Leu2251 | Asn–Leu | 3.60 |
| Leu2251 | Leu2251 | Leu–Leu | 4.00 | Leu2251 | Leu2251 | Leu–Leu | 3.37 |
| Leu2251 | Lys2252 | Leu–Lys | 3.98 | Lys2252 | Leu2251 | Lys–Leu | 3.81 |
| Thr2254 | Leu2255 | Thr–Leu | 3.94 | Leu2255 | Thr2254 | Leu–Thr | 3.90 |
| Lys2258 | Leu2255 | Lys–Leu | 3.66 | Lys2258 | Lys2258 | Lys–Lys | 3.65 |
| Lys2258 | Leu2262 | Lys–Leu | 3.83 | Asp2261 | Leu2262 | Asp–Leu | 3.51 |
| Leu2262 | Leu2262 | Leu–Leu | 3.73 | Leu2262 | Asp2261 | Leu–Asp | 3.90 |
| Gln2265 | Gln2265 | Gln–Gln | 3.99 | Gln2265 | Leu2269 | Gln–Leu | 3.70 |
| Leu2268 | Leu2269 | Leu–Leu | 3.75 | Leu2269 | Leu2269 | Leu–Leu | 3.75 |
| Leu2269 | Leu2272 | Leu–Leu | 3.97 | Leu2272 | Leu2269 | Leu–Leu | 3.88 |
| Leu2272 | Leu2272 | Leu–Leu | 3.92 | Leu2272 | Leu2276 | Leu–Leu | 3.78 |
| Lys2275 | Leu2276 | Lys–Leu | 3.71 | Leu2276 | Phe2279 | Leu–Phe | 3.78 |
| Phe2279 | Leu2280 | Phe–Leu | 3.77 | Leu2280 | Phe2279 | Leu–Phe | 3.65 |
| Asn2283 | Leu2286 | Asn–Leu | 3.76 | Glu2296 | Leu2297 | Glu–Leu | 3.54 |
| Leu2297 | Leu2297 | Leu–Leu | 3.47 | Leu2297 | Glu2296 | Leu–Glu | 3.96 |
| Gln2300 | Gln2300 | Gln–Gln | 3.46 | Lys2303 | Leu2304 | Lys–Leu | 3.87 |
| Leu2304 | Leu2304 | Leu–Leu | 3.46 | Met2307 | Leu2304 | Met–Leu | 3.76 |
| Leu2311 | Leu2311 | Leu–Leu | 3.67 | Leu2314 | Leu2311 | Leu–Leu | 3.83 |
| Leu2314 | Leu2314 | Leu–Leu | 3.96 | Asn2318 | Leu2321 | Asn–Leu | 3.53 |
| Leu2321 | Arg2322 | Leu–Arg | 3.63 | Arg2322 | Leu2321 | Arg–Leu | 3.65 |
| Glu2325 | Lys2328 | Glu–Lys | 3.96 | Lys2328 | Glu2325 | Lys–Glu | 3.86 |
| Lys2328 | Lys2328 | Lys–Lys | 3.82 | Ala2329 | Lys2328 | Ala–Lys | 3.93 |
| Glu2331 | Leu2332 | Glu–Leu | 3.80 | Glu2331 | Leu2332 | Glu–Leu | 3.85 |
| Leu2332 | Lys2328 | Leu–Lys | 3.93 | Leu2332 | Glu2331 | Leu–Glu | 3.61 |
| Leu2332 | Arg2335 | Leu–Arg | 3.73 | Arg2335 | Arg2335 | Arg–Arg | 3.97 |
| His2338 | Leu2339 | His–Leu | 3.69 | Met2342 | Val2346 | Met–Val | 3.81 |
| Lys2345 | Val2346 | Lys–Val | 3.62 | Met2349 | Val2353 | Met–Val | 3.88 |
| Leu2356 | Val2353 | Leu–Val | 3.93 | Leu2356 | Leu2356 | Leu–Leu | 3.81 |
| Leu2356 | Arg2357 | Leu–Arg | 3.65 | Val2360 | Val2360 | Val–Val | 3.96 |
| Lys2363 | Lys2363 | Lys–Lys | 3.64 | Leu2367 | Lys2363 | Leu–Lys | 3.68 |
| Leu2367 | Glu2366 | Leu–Glu | 3.79 | Gln2370 | Leu2367 | Gln–Leu | 3.88 |
| Gln2370 | Gln2370 | Gln–Gln | 3.83 | Lys2377 | Ile2381 | Lys–Ile | 3.62 |
| Ile2381 | Lys2377 | Ile–Lys | 3.65 | Ile2381 | Ile2381 | Ile–Ile | 3.86 |
| Ile2381 | Ile2381 | Ile–Ile | 3.51 | Ile2381 | Leu2384 | Ile–Leu | 3.96 |
| Leu2384 | Leu2384 | Leu–Leu | 3.80 | Leu2391 | Leu2391 | Leu–Leu | 3.86 |
| Glu2395 | Ala2394 | Glu–Ala | 3.91 | Lys2398 | Leu2402 | Lys–Leu | 3.84 |
| Ala2399 | Lys2398 | Ala–Lys | 3.93 | Glu2401 | Leu2402 | Glu–Leu | 3.65 |
| Leu2402 | Glu2401 | Leu–Glu | 3.70 | Leu2402 | Leu2402 | Leu–Leu | 3.75 |
| Leu2402 | Leu2405 | Leu–Leu | 3.78 | Leu2405 | Leu2402 | Leu–Leu | 3.79 |
| Leu2405 | Leu2405 | Leu–Leu | 3.85 | Leu2405 | Lys2406 | Leu–Lys | 3.85 |
| Val2409 | Val2409 | Val–Val | 3.62 | Glu2413 | Val2412 | Glu–Val | 3.51 |
| Leu2419 | Leu2419 | Leu–Leu | 3.81 | Leu2419 | Lys2420 | Leu–Lys | 3.49 |
| Leu2426 | Leu2426 | Leu–Leu | 3.98 | Asn2433 | Asn2433 | Asn–Asn | 3.96 |
| Glu2436 | Ile2437 | Glu–Ile | 3.81 | Ile2437 | Ile2437 | Ile–Ile | 3.62 |
| Leu2440 | Ile2437 | Leu–Ile | 3.97 | Leu2440 | Leu2440 | Leu–Leu | 3.87 |
| Ile2444 | Leu2440 | Ile–Leu | 3.94 | Val2447 | Ile2444 | Val–Ile | 3.64 |
| Val2447 | Val2447 | Val–Val | 3.83 | Val2447 | Ile2451 | Val–Ile | 3.56 |
| Ile2451 | Ile2451 | Ile–Ile | 3.80 | Ile2451 | Lys2450 | Ile–Lys | 3.51 |
| Ile2451 | Leu2454 | Ile–Leu | 3.90 | Leu2454 | Ile2451 | Leu–Ile | 3.86 |
| Leu2454 | Leu2454 | Leu–Leu | 3.78 | Leu2454 | Leu2458 | Leu–Leu | 3.77 |
| Leu2458 | Leu2454 | Leu–Leu | 3.63 | Leu2458 | Ile2461 | Leu–Ile | 3.97 |
| Ile2461 | Ile2461 | Ile–Ile | 3.67 | Arg2462 | Ile2461 | Arg–Ile | 3.48 |
| Leu2468 | Leu2468 | Leu–Leu | 3.83 | Leu2468 | Leu2468 | Leu–Leu | 3.99 |
| Arg2469 | Leu2468 | Arg–Leu | 3.89 | Pro2562 | Met2476 | Pro–Met | 3.86 |

Table S3: Hydrogen bonds at the FAZ10 central region dimer interface identified by PLIP.

| **Chain B** | **Chain A** | **Residues** | **Distance (Å)** |
|---|---|---|---|
| Asp2070 | Gln2165 | Asp–Gln | 2.79 |
| Ser2076 | Arg2166 | Ser–Arg | 3.75 |
| Ser2076 | Arg2166 | Ser–Arg | 3.75 |
| Asn2116 | Asp2070 | Asn–Asp | 3.08 |
| Tyr2121 | Arg2162 | Tyr–Arg | 3.02 |
| Arg2162 | Arg2072 | Arg–Arg | 3.38 |
| Arg2189 | Gln2184 | Arg–Gln | 3.64 |
| Lys2223 | Thr2224 | Lys–Thr | 3.05 |
| Thr2224 | Lys2223 | Thr–Lys | 2.73 |
| Arg2233 | Glu2238 | Arg–Glu | 4.00 |
| Glu2247 | Asn2248 | Glu–Asn | 3.05 |
| Asn2248 | Leu2244 | Asn–Leu | 3.32 |
| Lys2258 | Ser2259 | Lys–Ser | 3.31 |
| Ser2259 | Lys2258 | Ser–Lys | 3.53 |
| Gln2265 | Gln2265 | Gln–Gln | 2.73 |
| Gln2265 | Leu2262 | Gln–Leu | 2.88 |
| Asn2283 | Asn2283 | Asn–Asn | 2.79 |
| Asn2283 | Phe2279 | Asn–Phe | 3.07 |
| Ser2289 | Asp2290 | Ser–Asp | 2.60 |
| Asp2290 | Ser2289 | Asp–Ser | 2.65 |
| Asp2290 | Lys2293 | Asp–Lys | 3.21 |
| Lys2293 | Asp2290 | Lys–Asp | 2.92 |
| Gln2300 | Gln2300 | Gln–Gln | 2.89 |
| Glu2317 | Asn2318 | Glu–Asn | 2.86 |
| Asn2318 | Leu2314 | Asn–Leu | 3.83 |
| Arg2335 | Ser2336 | Arg–Ser | 3.02 |
| Ser2336 | Arg2335 | Ser–Arg | 3.62 |
| Ser2336 | Arg2335 | Ser–Arg | 2.83 |
| Gln2370 | Asn2371 | Gln–Asn | 3.52 |
| Asn2423 | Asn2423 | Asn–Asn | 2.96 |
| Asn2423 | Leu2419 | Asn–Leu | 2.97 |
| Asn2433 | Glu2430 | Asn–Glu | 2.85 |
| Ile2461 | Asn2465 | Ile–Asn | 3.49 |
| Asn2465 | Glu2464 | Asn–Glu | 3.00 |
| His2471 | Ser2473 | His–Ser | 3.41 |
| His2471 | Ser2473 | His–Ser | 3.49 |
| Ser2473 | His2471 | Ser–His | 3.06 |
| Ser2473 | His2471 | Ser–His | 2.88 |
| Arg2474 | Glu2467 | Arg–Glu | 3.90 |
| Ser2478 | Arg2564 | Ser–Arg | 3.64 |
| Tyr2523 | Arg2564 | Tyr–Arg | 2.92 |
| Arg2564 | Ser2473 | Arg–Ser | 2.92 |
| Arg2564 | Arg2521 | Arg–Arg | 3.05 |
| Arg2564 | Arg2521 | Arg–Arg | 3.72 |
| Gln2567 | Ser2478 | Gln–Ser | 2.76 |

Table S4: Salt bridges at the FAZ10 central region dimer interface identified by PLIP.

| Chain B | Chain A | Residues | Distance (Å) |
|---|---|---|---|
| Glu2078 | Arg2166 | Glu–Arg | 3.47 |
| Glu2105 | Arg2122 | Glu–Arg | 4.10 |
| Arg2119 | Asp2070 | Arg–Asp | 3.42 |
| Arg2122 | Glu2163 | Arg–Glu | 3.55 |
| Asp2123 | Arg2166 | Asp–Arg | 3.33 |
| Arg2155 | Asp2070 | Arg–Asp | 5.26 |
| Glu2163 | Arg2122 | Glu–Arg | 4.16 |
| Arg2166 | Asp2123 | Arg–Asp | 3.39 |
| Lys2170 | Asp2123 | Lys–Asp | 4.95 |
| Glu2212 | Arg2213 | Glu–Arg | 4.10 |
| Arg2213 | Glu2212 | Arg–Glu | 4.83 |
| Glu2247 | Lys2252 | Glu–Lys | 5.23 |
| Lys2252 | Glu2247 | Lys–Glu | 5.10 |
| Asp2290 | Lys2293 | Asp–Lys | 3.64 |
| Lys2293 | Asp2290 | Lys–Asp | 3.06 |
| Glu2317 | Arg2322 | Glu–Arg | 4.50 |
| Arg2322 | Glu2317 | Arg–Glu | 3.43 |
| Lys2328 | Glu2325 | Lys–Glu | 5.04 |
| Glu2352 | Arg2357 | Glu–Arg | 3.73 |
| Lys2377 | Asp2378 | Lys–Asp | 3.64 |
| Glu2413 | His2408 | Glu–His | 3.98 |
| Glu2422 | Arg2427 | Glu–Arg | 4.66 |
| Arg2427 | Glu2422 | Arg–Glu | 3.39 |
| Arg2469 | Glu2464 | Arg–Glu | 4.35 |
| Asp2472 | His2471 | Asp–His | 5.08 |
| Arg2474 | Glu2467 | Arg–Glu | 4.72 |
| Arg2524 | Glu2565 | Arg–Glu | 3.27 |
| Asp2525 | Arg2568 | Asp–Arg | 3.41 |
| Glu2565 | Arg2524 | Glu–Arg | 5.15 |
| Arg2568 | Asp2525 | Arg–Asp | 3.79 |

Table S5: $\pi$-cation interactions in the FAZ10 central region dimer interface identified by PLIP.

| Chain B | Chain A | Residues | Distance (Å) |
|---|---|---|---|
| Arg2474 | His2471 | Arg–His | 3.83 |
| Tyr2523 | Arg2564 | Tyr–Arg | 3.73 |

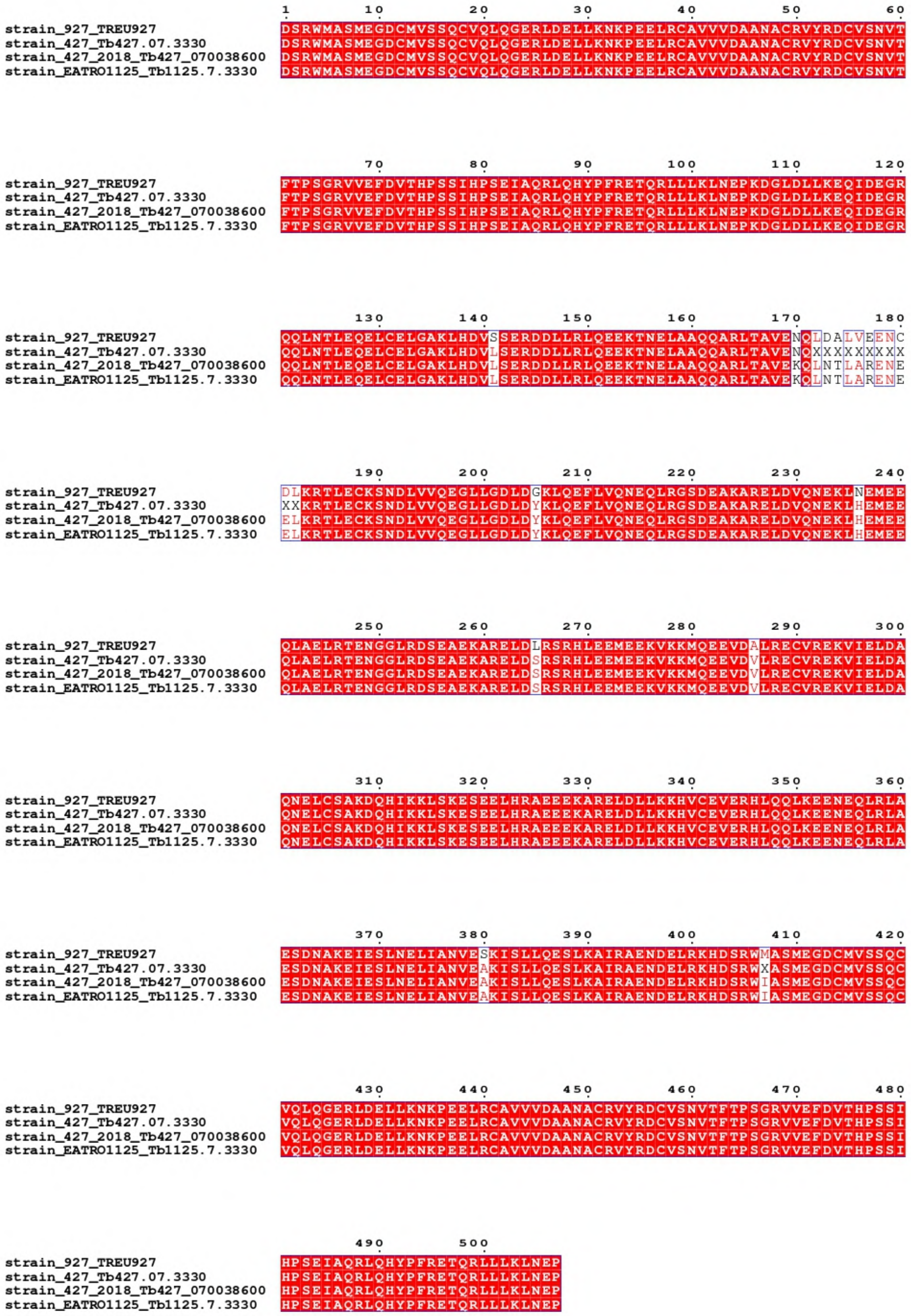


Figure S1: Sequence alignment of the FAZ10 central region across *Trypanosoma brucei* strains. Multiple sequence alignment of the FAZ10 central region from *T. brucei* strains 427, 427-2018, TREU927 and EATRO1125. Conserved residues are highlighted in red, with substitutions shown in white. The alignment was generated using Clustal Omega and visualized with ESPript 3.2. Positions annotated as "X" indicate residues that are not sequenced or undefined in the database. Amino acid numbering is shown relative to the central region (residues 1–507), corresponding to positions 2070–2576 in the full-length FAZ10 protein.

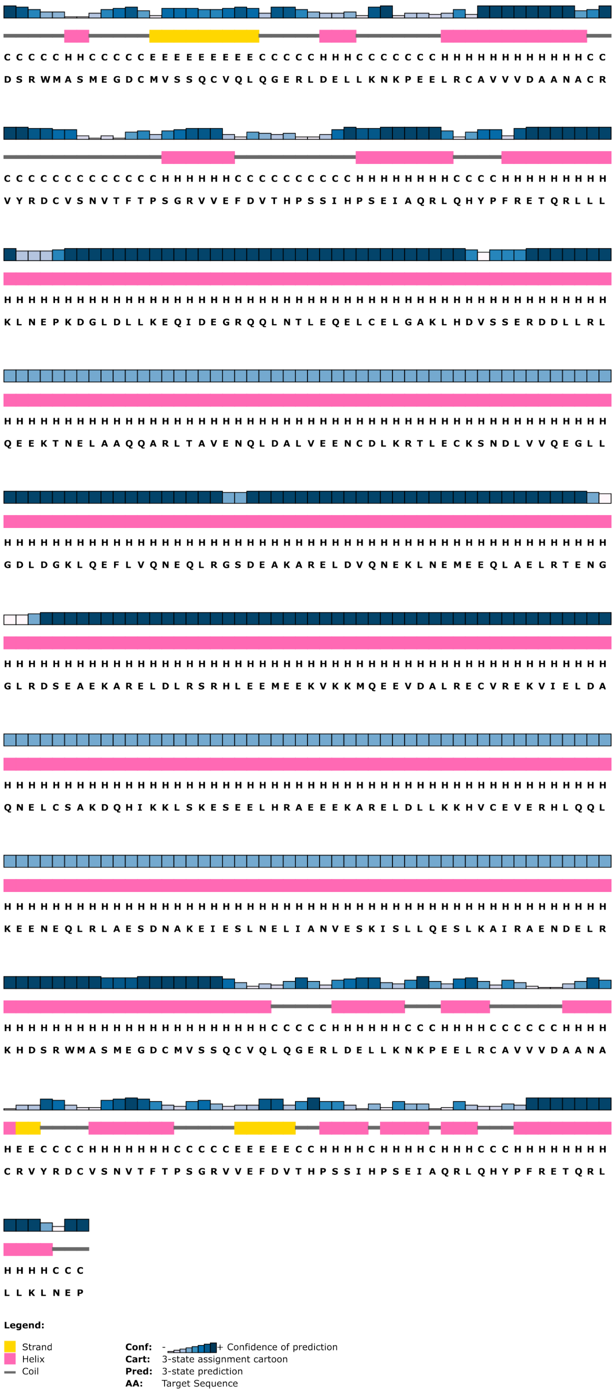


Figure S2: Secondary structure prediction of the FAZ10 central region using PSIPRED

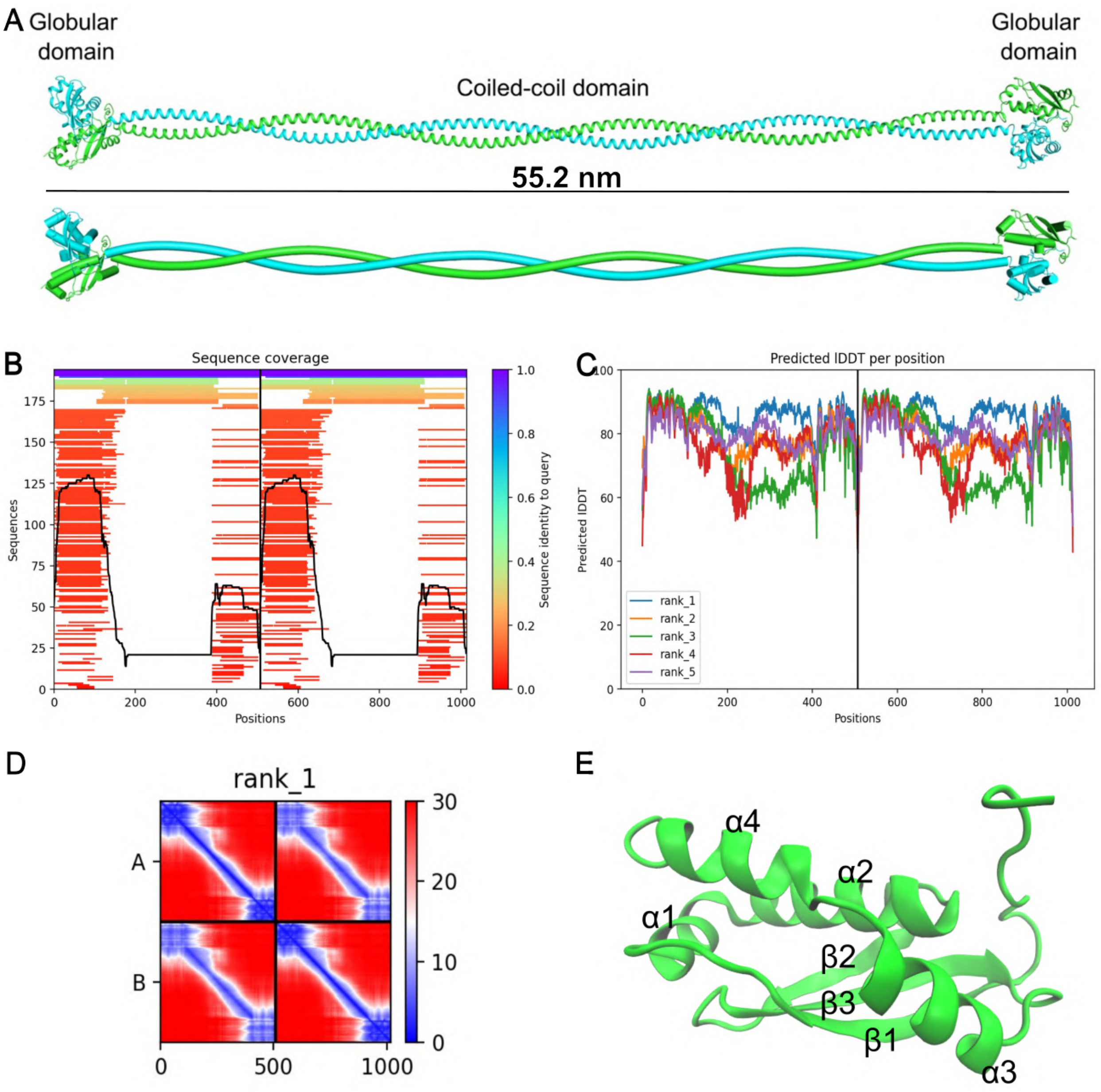


Figure S3: Structural prediction of the FAZ10 central region. (A) AlphaFold2 dimer model showing an elongated coiled-coil flanked by globular domains ( 44.2 nm). (B) Sequence coverage of the multiple sequence alignment used for modelling. (C) Per-residue confidence scores of the top-ranked model. (D) Predicted aligned error (PAE) map, showing low error within globular domains and higher uncertainty along the coiled-coil region. (E) Monomeric view of the terminal globular domain with $\alpha$-helices ($\alpha$1–$\alpha$5) and $\beta$-strands ($\beta$1–$\beta$3).

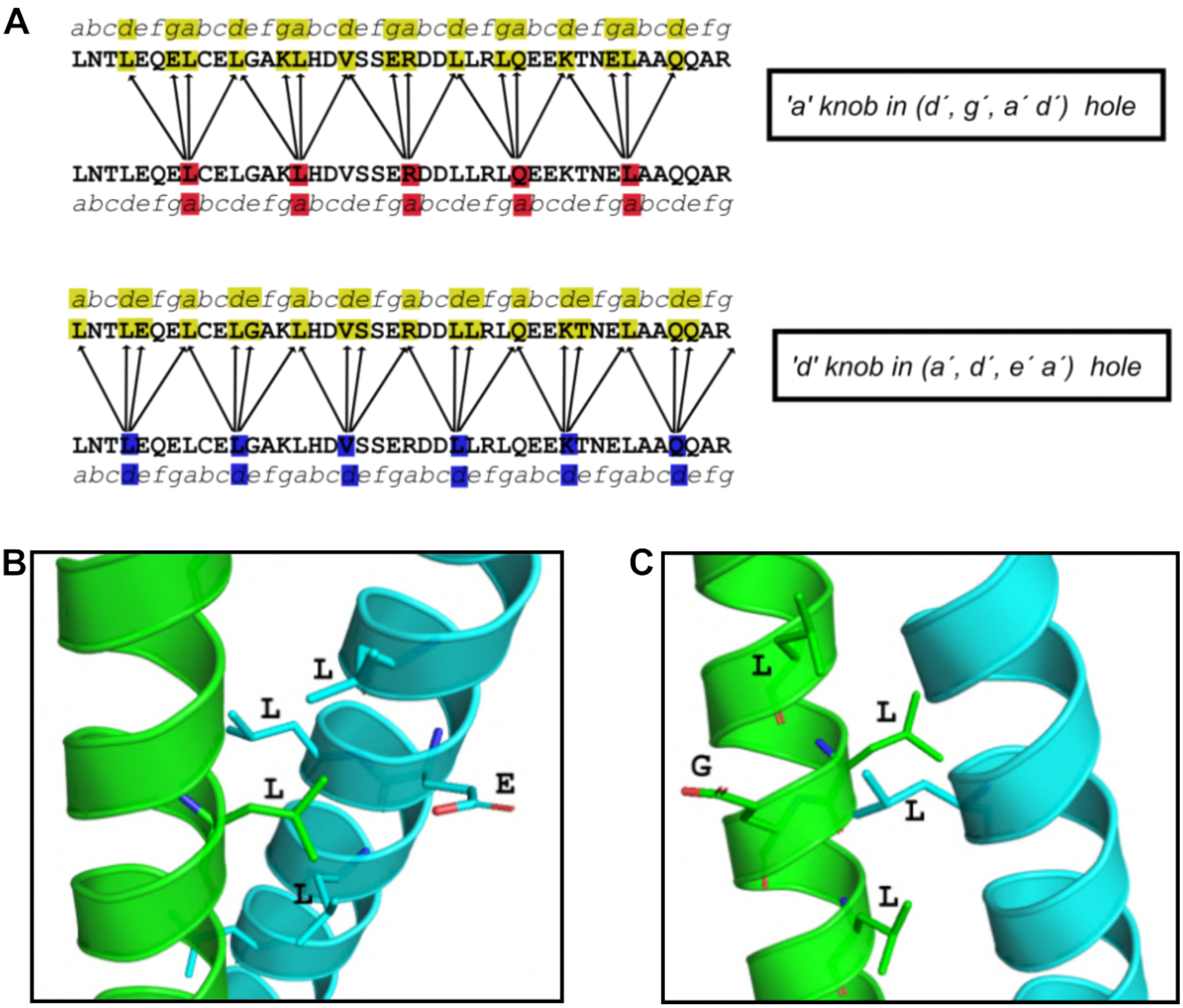


Figure S4: Structural and sequence-based characterization of the coiled-coil domain within the FAZ10 central region. (A) Coiled-coil dimer showing two parallel $\alpha$-helices. (B) Helical wheel diagram highlighting parallel chain orientation and canonical ‘a’/‘d’ heptad positions. (C) Schematic of predicted knobs-into-holes packing along the sequence. (D,E) Structural details of representative ‘a’ and ‘d’ knob–hole interactions, with interacting residues shown as sticks.

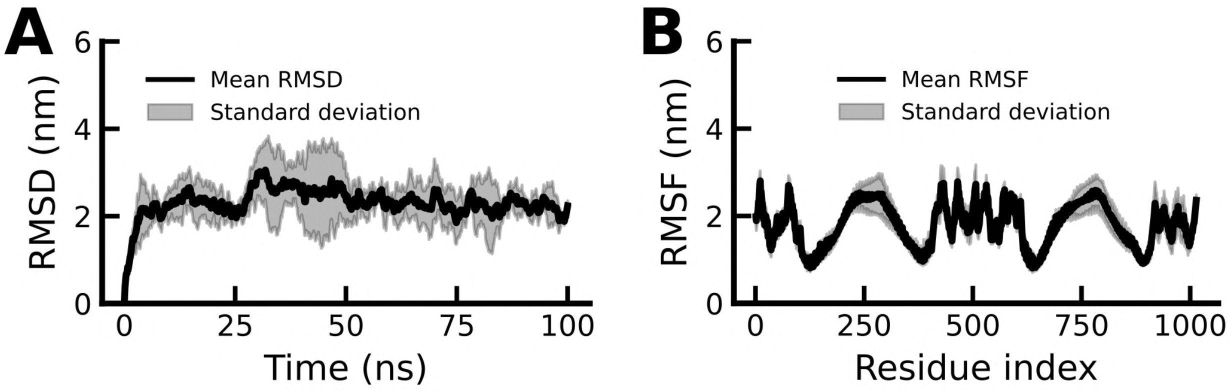


Figure S5: Stability of the FAZ10 central region assessed by all-atom molecular dynamics simulations (100 ns, three replicates). (A) Root mean square deviation (RMSD) and (B) root mean square fluctuation (RMSF) of the dimeric system.

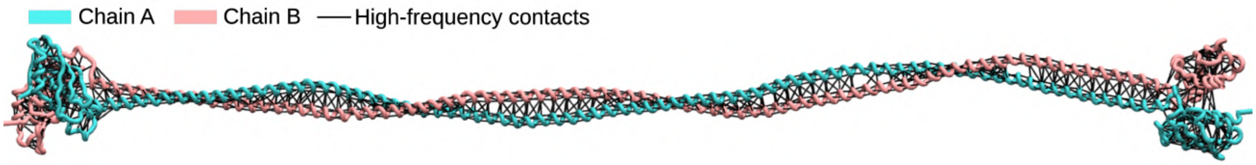


Figure S6: Spatial distribution of high-frequency contacts in the FAZ10 central region derived from all-atom molecular dynamics simulations (100 ns, three replicates).
